# Multiomics and quantitative modelling disentangle diet, host, and microbiota contributions to the host metabolome

**DOI:** 10.1101/2022.08.15.503927

**Authors:** Maria Zimmermann-Kogadeeva, Natasha A. Bencivenga-Barry, Michael Zimmermann, Peer Bork, Andrew L. Goodman

## Abstract

Dietary nutrients, host metabolism, and gut microbiota activity each influence the host’s metabolic phenotype; however, the interplay between these factors remains poorly understood. We employed tissue-resolved metabolomics in gnotobiotic mice carrying a synthetic human gut microbiota and germfree mice in two dietary conditions to develop an intestinal flux model that quantifies diet, host, and bacterial contributions to the levels of 2,700 intestinal metabolites. While diet was the main factor affecting metabolite profiles, we identified 1,117 potential microbial substrates and products in the gut. By integrating metagenomics and metatranscriptomics data into genome-scale enzymatic networks, we linked 202 potential substrate-product pairs by a single enzymatic reaction. We further identified bacterial species and enzymes that can explain the differential abundance of 13% of the identified microbial products between the mouse groups. This quantitative modelling approach paves the way for controlling an individual’s metabolic phenotype by modulating their gut microbiome composition and diet.

## Introduction

The gut microbiota comprises hundreds of microbial species residing along the intestine^1^, which come into contact with dietary and host-produced compounds that transit through the gastrointestinal tract during digestion. Although diet rapidly and reproducibly changes microbiota composition^2,3^, many microbial species are repeatedly detected in the gut for months, years and decades, indicating that microbes can adapt their metabolism to changing nutrient environments^4–8^. Some metabolic changes are restricted to the gut, with potential impacts on other microbes and gut epithelial cells^9,10^. Other microbe-derived metabolites are absorbed from the intestine and distributed systemically through blood circulation so that they can be detected in serum^11,12^ and peripheral organs^13^, such as liver and brain^14–16^. Microbiota-produced metabolites span a broad range of chemical classes^17,18^, including amino acids and peptides, fatty acids and lipids^9,19^, glycans^20,21^, simple and complex polysaccharides^22^, bile acids^23^, hormones and neurotransmitters^14^. Metabolites of different origin across tissues collectively define a host metabolic phenotype that can impact diverse aspects of health and disease risk.

Understanding the diet-microbiome relationship is essential for efforts to design dietary, microbial, or other interventions aimed at a desired beneficial metabolic phenotype. With the increasing availability of microbiome and metabolome data in humans, it becomes possible to associate metabolite variation to microbial abundance^24–26^. However, mechanistic insights into microbe-metabolite links are challenged by many confounding factors, including variations in host genotype, diet, environmental exposure, and others^27,28^. Although it is in principle possible to modulate the gastrointestinal and circulating metabolite pools through diet and microbiome interventions^29–32^, it remains challenging to predict these interactions because the contributions of diet, host, and microbiota to metabolite levels remains mostly undefined.

Here we combine gnotobiotic mouse experiments with dietary interventions and multiomics data integration to disentangle diet, microbiota and host contributions to the metabolic profiles in the host. Using six complementary mass spectrometry protocols, we measured 57,340 metabolic features corresponding to 4,649 annotated metabolites and classified them based on their distribution profiles across tissues. We developed a coarse-grained intestinal metabolic flux model, which quantifies diet, host and microbiome contributions to metabolite abundance in the gastrointestinal tract (GIT). We further explore metagenome-wide enzymatic reaction networks to identify candidate metabolite substrate-product pairs and bacterial enzymes potentially responsible for their conversion. This approach for integrating spatial metabolomic measurements with quantitative mathematical modelling and multiomics integration can be applied across diets, perturbations, host genetic backgrounds and microbiome compositions to ultimately identify decisive factors affecting metabolite levels and inform rational design of microbiota therapies.

## Results

### Diet influences both microbiota composition and gene expression

To disentangle relationships between diet, host and microbiota, we conducted a controlled diet and microbiota experiment in a gnotobiotic mouse model. We colonized germfree C57BL/6 mice with a synthetic, defined community of 18 genome-sequenced human gut bacteria that represent four major phyla common in the human gut and were previously successfully used in synthetic communities^29,33^. This community encodes a diverse metabolic potential, including breakdown of complex polysaccharides, amino acid fermentation, and acetogenesis^29,34^ (Figure 1a, Supplementary Table 1). Germfree and colonized mice were fed either a high-fat diet (HFD) or a calorie-matched high-carbohydrate diet (HCD) for four weeks (Figure 1a, Supplementary Figure 1a-c, Supplementary Table 2), at which point we collected samples along the gastrointestinal tract (GIT), serum and liver for metabolomics analysis, and cecal content of colonized mice for metagenomic and metatranscriptomic sequencing (Figure 1a).

**Figure 1.**
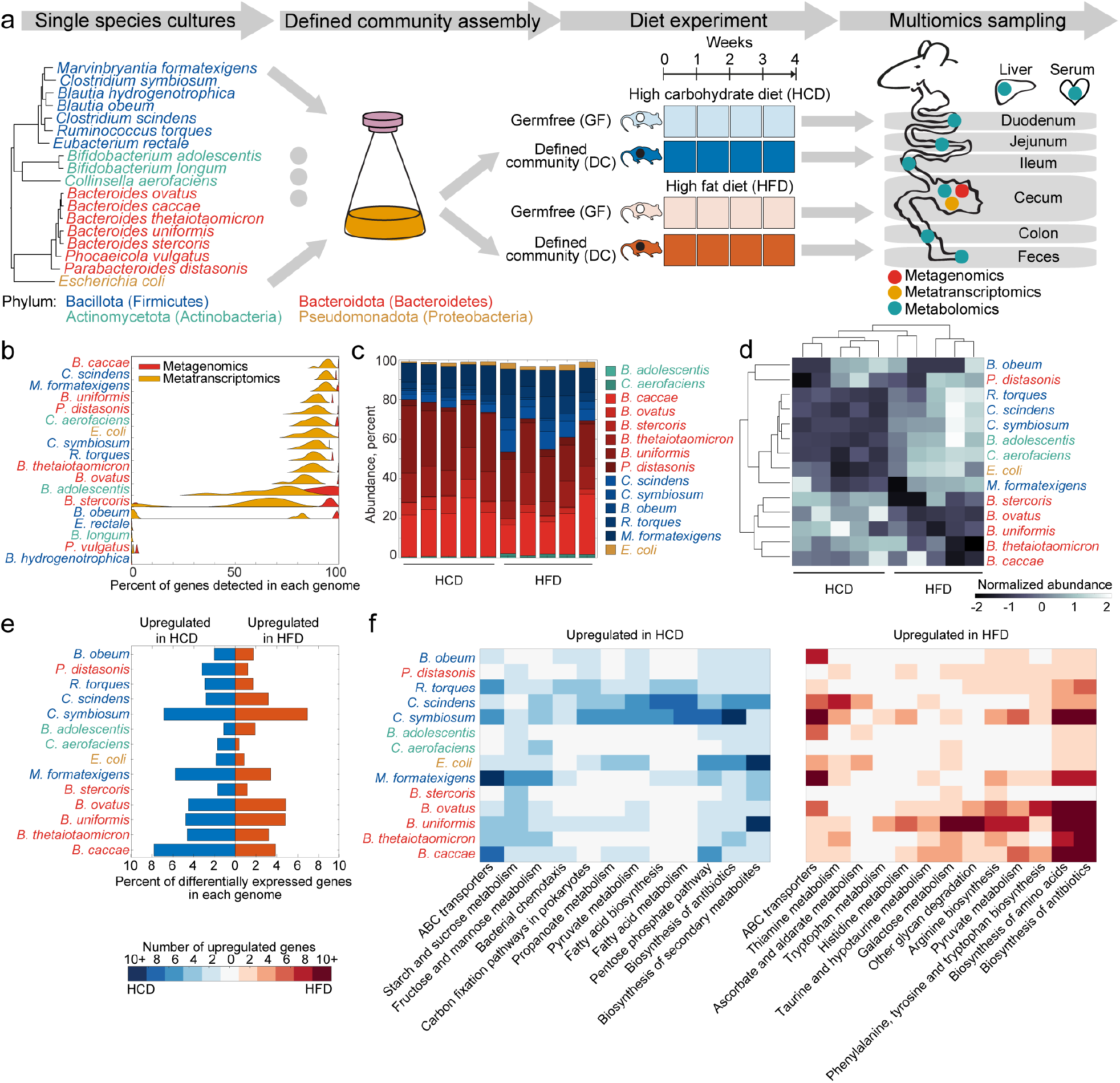
Controlled gnotobiotic mouse experiment to capture diet adaptation in a synthetic gut microbiota. **a.** Schematic representation of the diet experiment and sample collection. **b.** Distribution of the percent of detected genes per species in metagenomics and metatranscriptomic measurements across all colonized mouse samples. **c.** Stacked barplot representing relative species abundance estimated from metagenomics data across individual animals (n=10). **d.** Hierarchical clustering of normalized species abundances across individual animals (n=10). **e.** Percent of genes differentially expressed between the two diet conditions in each detected species (abs(log2(fold change)) ≥ log2(1.5), FDR-adjusted p-value ≤ 0.05). **f.** Pathway enrichment analysis of genes differentially expressed by each species in each of the diet conditions. Only significantly enriched pathways (FDR-adjusted p-value ≤ 0.1) with at least three differentially expressed genes in at least one species are depicted.

The microbial load was comparable between all colonized animals consuming either of the two diets (Supplementary Figure 2a). Metagenomic and metatranscriptomic sequencing reached an average depth of 2.5 mio reads (~750 Mb) and 5 mio reads (~1,500 Mb) across samples, respectively. This allowed us to stably recover reads unique to 14 of the 18 species from the defined community with a high coverage of bacterial genomes (average of 90% (metagenomics) and 80% (metatranscriptomics) of genes); missing species were below detection at our limit of resolution (<0.25%, Figure 1b), and were likely outcompeted by the other community members. We also recovered sequences that mapped to *Lactococcus lactis*, which was not included in the defined community but is known to be present but nonviable in autoclaved casein-rich diets^35^ (Supplementary Table 3). Principal component analysis (PCA) of metagenomic and metatranscriptomic datasets revealed consistent grouping of the samples by the administered diet (Supplementary Figure 2b, c), suggesting diet-dependent signatures of microbial abundances and transcriptional activity.

Representatives of the phylum Bacteroidota (formerly Bacteroidetes) dominated the microbiome in both colonized mouse groups, and their relative abundance decreased on HFD with concordant increase in the abundance of Bacillota (formerly Firmicutes), Actinomycetota (formerly Actinobacteria) and Pseudomonadota (formerly Proteobacteria) (Figure 1c). Hierarchical clustering of relative species abundances normalized across all samples separated mice from the two diet groups, and separated bacteria according to their phylogenetic relationships, in concordance with previous reports that Bacteroidota thrive in high-carbohydrate conditions, and Bacillota, Actinomycetota and Pseodomonadota are often more abundant on HFD^35^ (Figure 1d). There was a general agreement between relative DNA and RNA abundance per species and sample (Pearson’s correlation coefficient, PCC = 0.43), although the relative RNA species abundance and the RNA/DNA ratio was higher for some Bacillota compared to other species (Supplementary Figure 2d-h).

Taken together, our controlled diet and microbiota experiment provides high coverage data for microbiota abundance and gene expression, and establishes that previously reported diet-specific changes in microbiota composition are recapitulated in this model system.

### Diet induces transcriptional changes in all bacterial species in the community

Differential analysis of transcriptional profiles of the 14 stably detected species revealed that all species changed their gene expression between the two diets, with some species differentially expressing up to 10% of their genes (abs(log2(fold change)) ≥ log2(1.5), FDR ≤ 0.05, Figure 1e, Supplementary Table 4). Pathway enrichment analysis of differentially expressed genes reveals patterns of common and phylum-specific adaptation of bacteria to either of the diets. For example, starch is the main ingredient of the HCD; representatives of all phyla upregulated genes involved in starch and sucrose, fructose and mannose metabolism, and the pentose phosphate pathway on this diet (Figure 1f, Supplementary Table 5). Fatty acid biosynthesis and metabolism genes were upregulated on HCD compared to HFD by representatives of Bacillota, suggesting that on HCD, these species need to synthesize fatty acids that they may be able to obtain directly from the HFD. This hypothesis is supported by previous reports that representatives of Bacillota, and specifically Clostridiaceae, are capable of uptake and utilization of long-chain fatty acids^36,37^. On HFD, most species upregulated amino acid biosynthesis pathways, possibly indicating that amino acids are less available for uptake in HFD compared to HCD. Most species also upregulated genes involved in thiamine metabolism (Figure 1f), which might reflect the increased need to synthesize thiamine on HFD compared to HCD. Notably, thiamine is supplemented to the latter diet (Supplementary Figure 1c). Consistent with previous reports^38^, we observed Bacteroidota-specific regulation of glycan degradation on HFD, suggesting that these bacteria shift from plant glycan to mucosal glycan foraging on HFD. Together, these results demonstrate an orchestrated general microbial response to the two different diets, and point out some phyla-specific adaptations linked to fatty acid, polyketide sugar unit or glycan metabolism.

### Levels of metabolites in the host are tissue-, diet- and microbiota-dependent

Next, we investigated the impact of diet and microbiota on metabolite levels across different tissues, employing six complementary protocols for untargeted metabolomics analysis. Altogether, we detected 57,340 unique metabolic features, 4,649 of which we could annotate using the Kyoto Encyclopedia of Genes and Genomes (KEGG)^39^ and the Human Metabolome Database (HMDB)^40^ metabolite repositories (Materials and Methods, Supplementary Table 6). In each tissue, between 1,242 and 2,176 metabolic features were annotated, and the majority of metabolites were detected both in germfree and colonized mice (Figure 2a). Between 28% and 78% of annotated metabolites overlapped between tissues (Supplementary Figure 3a). PCA applied to all metabolomics samples revealed grouping by tissue, with serum and liver metabolite profiles being distinct from the GIT tissues, which grouped according to physiological proximity (Figure 2b). PCA of individual tissues revealed grouping based on diet and mouse colonization state in serum and the three sections of the large intestine; in liver and the small intestine, samples from different conditions were more similar to each other (Figure 2b, Supplementary Figure 3b).

**Figure 2.**
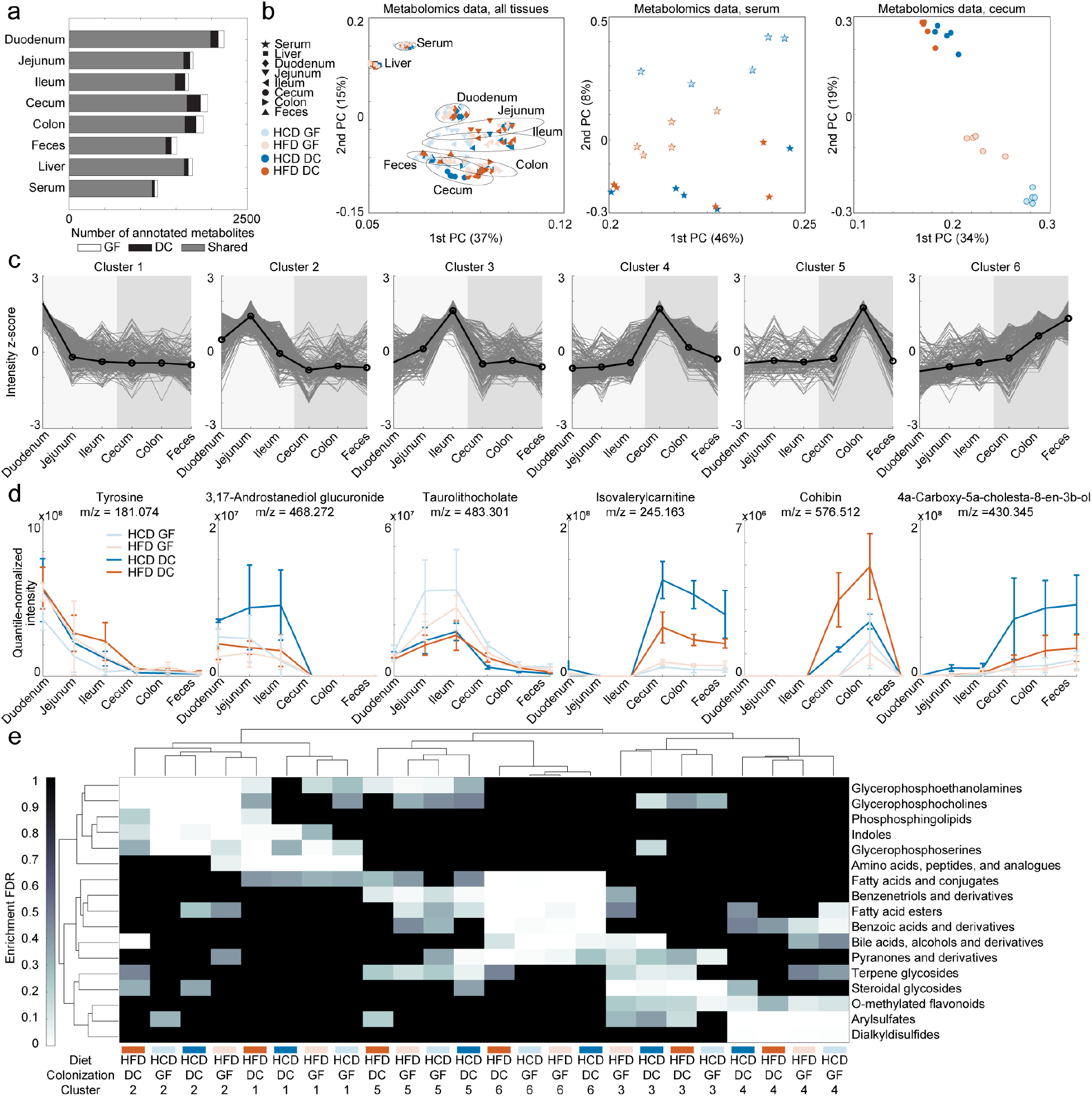
Spatial distribution of metabolites across tissues is consistent across diets and colonization states. **a.** Number of annotated metabolites per mouse tissue. **b.** PCA of annotated metabolites in all tissues (left), serum (center) or cecum (right). **c.** K-means clusters of normalized metabolite profiles in the GIT of germfree mice consuming HCD. **d.** Examples of metabolites belonging to each of the six clusters in each of the four mouse groups. **e.** Hierarchical clustering of the results of chemical group enrichment analysis of the metabolites belonging to each of the k-means clusters. Analysis was performed separately for each of the four mouse groups. Only significantly enriched groups (FDR-adjusted p-value ≤ 0.1) are depicted. HFD – high-fat diet, HCD – high-carbohydrate diet, GF – germ-free mice, DC – mice colonized with the defined community.

### Spatial profiles of metabolites across tissues are consistent across diet- and microbiota-conditions

To identify sources of different metabolites, we next set out to investigate tissue-specific metabolite profiles starting with the GIT. For each experimental condition, we normalized metabolite intensities across the six sections of the GIT and performed unsupervised k-means clustering of the normalized metabolite profiles. For each of the four conditions, the clustering revealed six distinct clusters characterized by peak metabolite intensity in one of the six sections of the GIT (Figure 2c, Supplementary Figure 4a, b). These clusters are linked to host physiology; for example, the amino acid tyrosine is assigned to Cluster 1 with peak intensity in duodenum, consistent with its absorption from the small intestine as a dietary nutrient (Figure 2d). Androstanediol glucuronide, assigned to Cluster 2 with peak intensity in jejunum, and taurolitocholate, assigned to Cluster 3 with peak intensity in ileum, are known host metabolites produced in the liver, and are secreted to the small intestine via the biliary duct. A carnitine derivative isovalerylcarnitine assigned to Cluster 4, a fatty acyl cohibin assigned to Cluster 5, and cholesterol derivative 4a-Carboxy-5a-cholesta-8-en-3b-ol assigned to Cluster 6, with peak intensities in cecum, colon and feces, respectively, likely reflect the propagation of native and diet-derived compounds through the large intestine and subsequent secretion via feces (Figure 2d). These spatial distributions are in concordance with current knowledge on metabolic aspects of the digestive process.

To systematically characterize metabolites belonging to each cluster in each condition, we performed HMDB chemical class enrichment analysis. Hierarchical clustering of the enriched chemical classes revealed similarities of their tissue distributions independent of diet or mouse colonization state (Figure 2e). Thus, Cluster 1 metabolites belong to the classes of amino acids, fatty acids, glycerophosphoserines and indoles, and are likely dietary nutrients absorbed in duodenum. Clusters 2 and 3 contain phosphosphingolipids, glycerophosphocholines, bile acids, and steroidal glycosides, that are likely produced in the liver and gallbladder and secreted into the small intestine via bile to aid digestion and excretion from the body. Clusters of metabolites with peak intensities in the large intestine contain various classes of fatty acids and fatty acid esters, bile acids, benzenetriols, benzoic acids, pyranones, arylsulfates and dialkyldisulfides, potentially representing the results of host digestion of dietary nutrients and compounds produced by the host along the GIT (Figure 2e).

Metabolites detected in serum and liver were highly overlapping: 833 metabolites were detected in both tissues, corresponding to 67% of serum and 48% of liver metabolites (Supplementary Figure 3a). By contrast, serum and liver samples shared between 49% and 37% of metabolites with GIT samples, with the number of shared metabolites decreasing from ileum to feces as expected due to the decrease in intestinal uptake, and thus exchange of metabolites between intestinal lumen and serum, from ileum to feces. Metabolites belonging to chemical classes of glycerophosphoethanolamines and terpene glycosides were shared between liver and serum, but not GIT, while o-methylated flavonoids, steroidal glycosides, and glycerophosphoserines were either serum- or liver-specific independent of diet or mouse colonization state (Supplementary Figure 4c). Together, these results demonstrate that metabolite profiles across tissues reflect expected host physiology, and that the experimental parameters (germfree or defined community, HCD or HFD) do not globally disrupt the spatial distribution of metabolites in host tissues.

### Development of intestinal flux model to assess sources of metabolites in the GIT

While tissue distributions of metabolites were consistent across the four mouse groups, we next set out to compare metabolite levels in each tissue across conditions. Both diet and microbiota colonization affected the levels of several hundred metabolites, with the largest number of differentially abundant metabolites detected in the large intestine (Supplementary Figure 5, Supplementary Table 7). For example, while the cluster assignments of glutamate, propionate and L-octanoylcarnitine were consistent across mouse groups (Cluster 1, Cluster 6 and Cluster 4, respectively), the levels of glutamate and L-octanoylcarnitine were significantly higher in the large intestine of germfree mice, whereas the levels of propionate were significantly higher in the large intestine of colonized mice (FDR-adjusted p-value < 0.03, Figure 3a, Supplementary Table 7).

**Figure 3.**
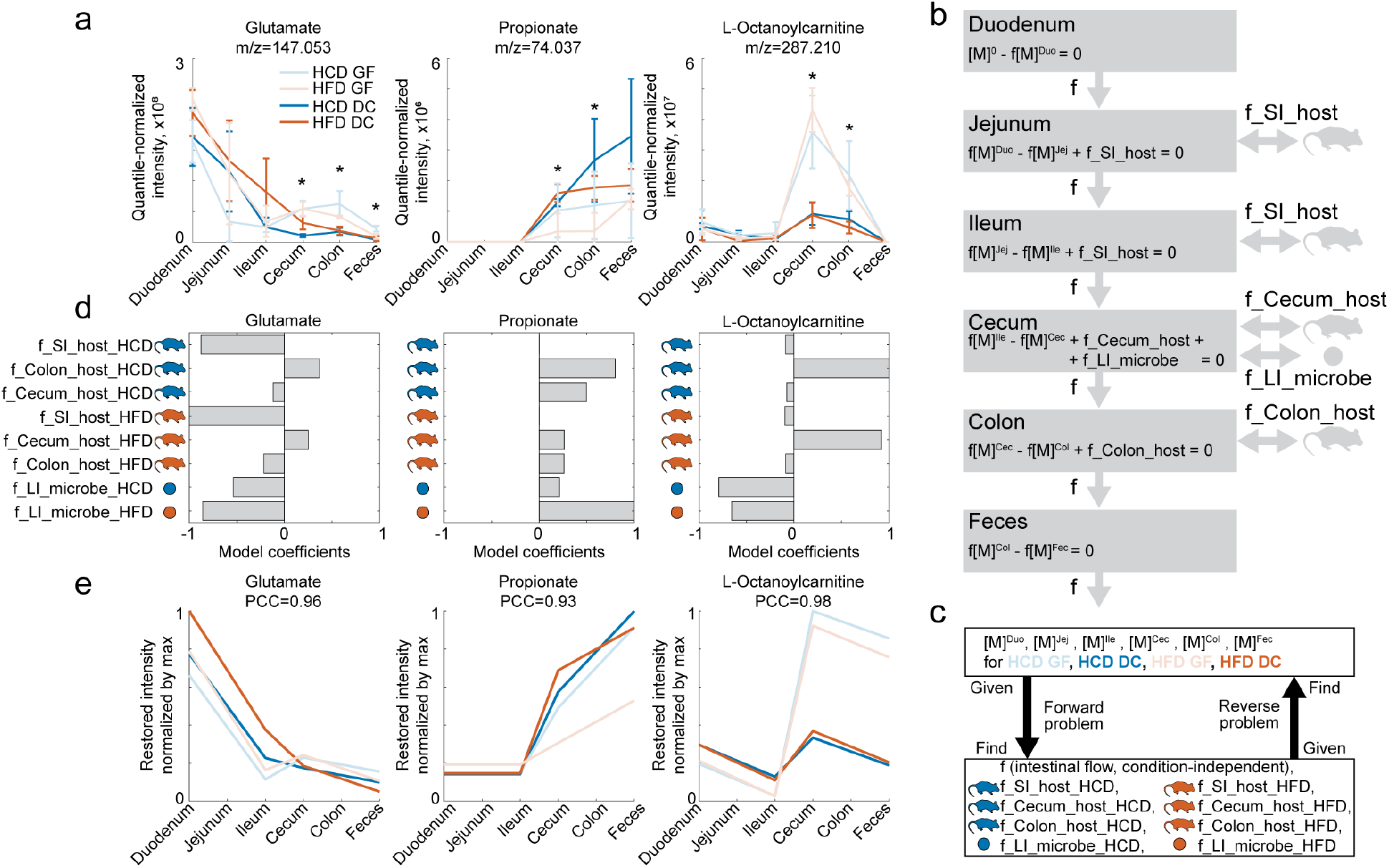
Coarse-grained intestinal flux model estimates diet, host and microbiome contributions to the metabolite profiles in the GIT. **a.** Example profiles of three metabolites detected in the GIT in the four conditions. * indicate FDR-corrected p-value < 0.05 (two-sided t-test) for comparison between GF and DC mouse groups in at least one of the diet conditions. **b.** Scheme of the coarse-grained intestinal flux model. **c.** Scheme of inputs and outputs of the forward and reverse problems aimed at estimating model parameters and restoring metabolic profiles, respectively. **d.** Host and microbiome metabolism coefficients in HCD and HFD estimated by the model for each of the metabolites in (a), normalized by the largest value. **e.** Restored metabolite profiles obtained by solving the reverse problem (c) for each metabolite in (a) and set of parameters in (d). PCC indicate Pearson’s correlation coefficient between restored (e) and original (a) metabolite profiles. HFD – high-fat diet, HCD – high-carbohydrate diet, GF – germ-free mice, DC – mice colonized with the defined community.

Because metabolite profiles were generally consistent with expected host physiology, we reasoned that they should reflect diet, host, and microbiota contributions to metabolite levels in the GIT, and set out to establish a coarse-grained intestinal flux model that describes these processes. In each GIT compartment, we described the changes in metabolite levels with an equation including compartment-specific consuming and producing metabolic fluxes. The model connects GIT compartments via intestinal flux which represents metabolite transit. In each compartment, the change in metabolite abundance due to intestinal transit follows the law of mass action with a condition- and compartment-independent parameter f (Figure 3b). In the small intestine, parameter f_SI_host describes host metabolism flux, and in the large intestine host metabolism fluxes are described by parameters f_Cecum_host and f_Colon_host for cecum and colon, respectively. In the cecum of colonized mice, metabolite levels are also influenced by microbiota metabolism flux parameter f_LI_microbe (Figure 3b). In this simple model, we assumed that host metabolism parameters are independent of mouse colonization state, but treated all host and microbiota metabolism parameters as diet-specific. The resulting model contained 20 equations (for jejunum, ileum, cecum, colon and feces for each of the four conditions) with nine parameters (the nonnegative intestinal flux parameter f and eight diet-specific host and microbiota metabolism parameters which can have either negative or positive values corresponding to metabolite consumption or production; Figure 3c, Supplementary Table 8).

Since the spatial metabolite profiles were highly similar across these *ad libitum-fed* animals, we assumed that the measurements reflect a pseudo-steady state, *i.e*., metabolite concentrations in each GIT compartment are stable over time. Under this assumption, the nine model parameters can be estimated in a so-called “forward problem”: a system of linear equations, where metabolite levels in each tissue and each condition were used as coefficients for the intestinal flow parameter f, and coefficients of all other parameters were equal to one (Figure 3c, Materials and methods). To assess the quality of the model, we formulated the so-called “reverse problem”: in the same system of linear equations, the intestinal flux parameter and host and microbiota metabolism parameters estimated in the forward problem are used as coefficients, and the system is solved to estimate metabolite levels in each tissue and each condition (Figure 3c, Supplementary Table 8). We then correlate the restored metabolite profiles across conditions with the original data to assess how well the estimated parameters in the model recapitulate the measured metabolic profiles in the GIT. For further analysis, we consider metabolites for which the restored metabolite intensities calculated by solving the reverse problem correlate with the original metabolite profiles at PCC ≥ 0.7.

### Intestinal flux model distinguishes sources of metabolite abundance in the GIT

To quantify the relative contributions of diet, host and microbiota to a metabolite profile in the GIT, we estimated the model parameters and normalized the values of the eight condition-dependent metabolic flux parameters to the largest absolute value. While we did not explicitly include diet as a factor in our model, we reasoned that metabolites whose primary source is diet are characterized by large negative host metabolism parameters in the small intestine (f_SI_host), corresponding to host consumption or absorption in this tissue. Metabolites most influenced by host production are characterized by large positive host metabolism parameters in the small and/or large intestine (f_SI_host, f_Cecum_host, f_Colon_host), whereas metabolites whose levels are affected by microbiota production will have large positive microbiota metabolism parameter values (f_LI_microbe).

To test this approach, we first examined metabolites whose sources are known. For example, the major contributors to the profile of the amino acid glutamate are the negative host metabolism parameters in the small intestine (Figure 3d), suggesting that diet is the main source of glutamate in the GIT. The model also estimates negative bacterial metabolism parameters in the large intestine, suggesting that bacteria consume residual glutamate from the GIT. Propionate provides a second example: according to the model, its profile is mainly affected by bacterial production (positive bacterial metabolism parameter on both diets (Figure 3d)), consistent with reported production of propionate by the gut microbiota^41–43^. In a third example, the profile of L-octanoylcarnitine is influenced by host production in the large intestine, consistent with it being a product of fatty acid oxidation by the host. Similar to glutamate, negative microbial metabolism parameters indicate that bacteria consume this metabolite in the large intestine (Figure 3d). For each of these metabolites, metabolite profiles restored by solving the reverse problem highly correlated with the original metabolite measurements (PCC ≥ 0.93, Figure 3e).

Together, these results demonstrate that the coarse-grained intestinal flux model can disentangle relative contributions of diet, host and microbiome to the metabolite profiles, which we next set out to systematically assess for all metabolites detected in the GIT.

### Diet is the major factor affecting metabolite profiles in the GIT, followed by host and microbiota metabolism

We applied the intestinal flux modelling approach to all 3,716 annotated metabolites detected in the GIT. For 2,700 (73%) of these metabolites, the identified parameters could reproduce measured metabolite levels in the reverse problem (PCC ≥ 0.7). Random shuffling of these parameters removed this correlation (Figure 4a).

**Figure 4.**
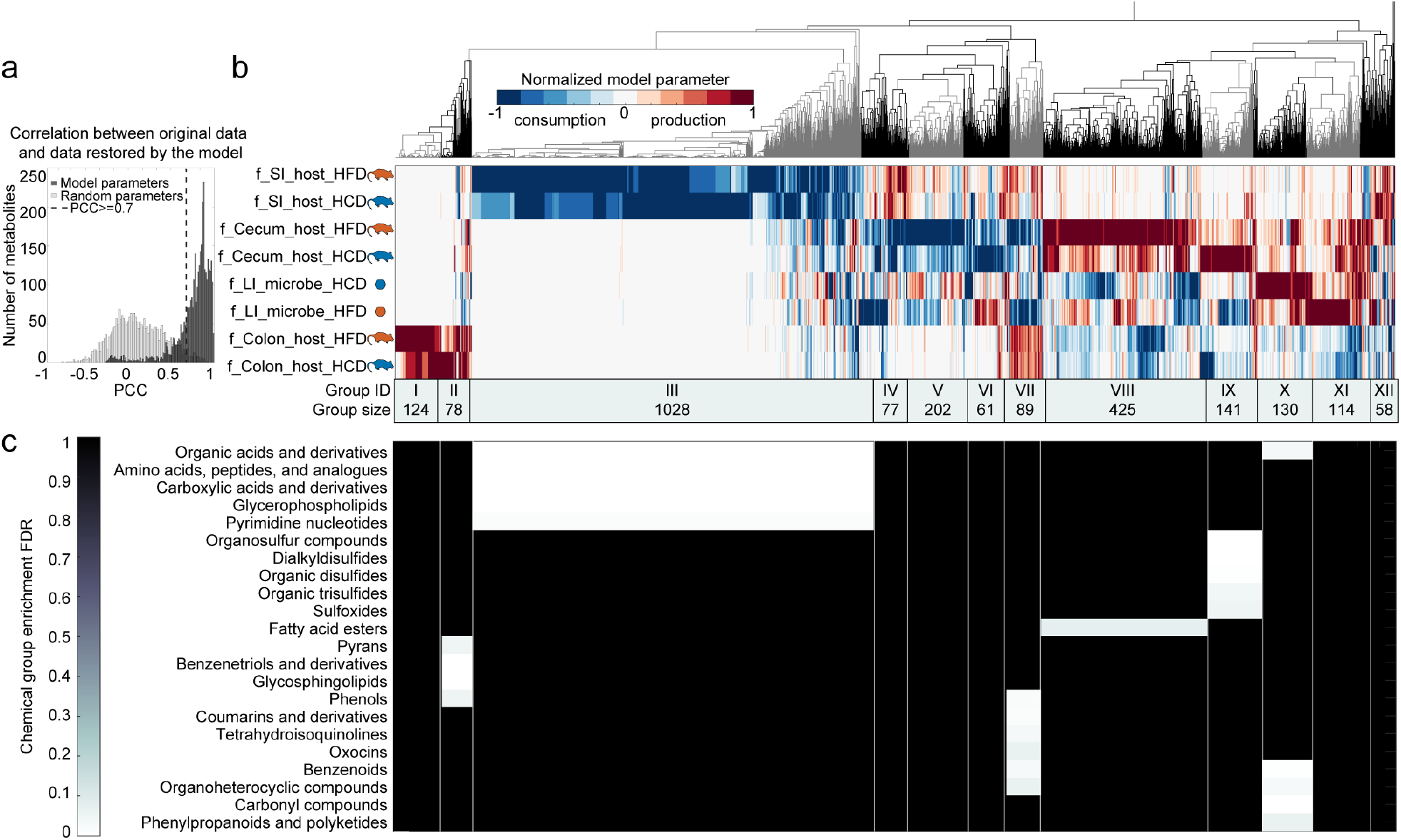
Diet contributes most to the metabolite profiles in the GIT, followed by host and microbial metabolism. **a.** Distributions of correlation coefficients between the original metabolite profiles and restored profiles obtained by solving the reverse problem with either estimated model parameters or randomly shuffled model parameters. **b.** Hierarchical clustering of normalized model parameters estimated for 2,700 metabolites stably detected in the GIT, for which profiles restored in the reverse problem closely correlated with the original data (PCC ≥ 0.7). Grey and black colors of the dendrogram branches indicate manually defined groups of metabolites with similar normalized profiles of estimated model parameters. **c.** Chemical group enrichment results of metabolites belonging to the groups manually defined in the clustergram in (b). Only significantly enriched chemical groups (FDR-adjusted p-value ≤ 0.1) are depicted.

To get an overview of the predicted factors contributing to each of the metabolite profiles, we performed hierarchical clustering of the normalized values of the eight metabolic flux parameters, which allowed us to define 12 distinct metabolite groups characterized by the main contributing parameter (Figure 4b). Thus, Groups I and II contain metabolites with a high host metabolism parameter in the colon on HFD or HCD, respectively; Group III contains metabolites with a high negative host metabolism parameter in the small intestine on both diets; whereas Groups X and XI contain metabolites with a high positive microbial metabolism parameter on HCD and HFD (Figure 4b). Group III characterized by the negative host metabolism parameter in the small intestine is the largest of these groups, containing 38% (1,028/2,700) of the modelled metabolites; this suggests that diet is the major factor affecting metabolite profiles in the GIT. About one third of the analysed metabolites are potentially produced by the host, since they belong to the clusters with high host metabolism parameters in the small intestine (Groups IV and XII, in total 5%, or 135/2,700 metabolites), or in the large intestine (Groups I, II, VIII and IX, in total 28%, or 768/2,700 metabolites). We found that 9% of metabolites (244/2,700) belong to the clusters representing models in which microbial metabolism was the highest contributing parameter (Groups X and XI), suggesting that these metabolites are produced by (or in response to) gut bacteria.

To systematically characterize metabolites affected by different factors, we performed HMDB chemical class enrichment analysis for each group (Figure 4c, Supplementary Table 7). Amino acids, organic acids, carboxylic acids and glycerophospholipids were enriched in Group III (diet-derived metabolites). Glycosphingolipids, phenols, benzenetriols and pyrans were enriched in clusters defined by positive host colon metabolism parameters, consistent with host production. Other enrichments are diet-specific: for example, fatty acid esters were enriched in Group VIII (positive cecum host metabolism parameter on HFD), whereas organic sulfides and sulfoxides were enriched in Group IX (positive cecum host metabolism parameter on HCD), consistent with expected host metabolism products of the corresponding diet ingredients. Metabolites from Group X (characterized by a high microbial metabolism parameter) were enriched in organic acids, benzenoids, carbonyl compounds and polyketides, which is in line with prior reports of microbiota-associated compounds^44–47^.

This approach focuses on the largest model-identified parameter to highlight primary factors contributing to a metabolite profile in the GIT. However, we observed that for many metabolites (including glutamate and L-octanoylcarnitine), for which host parameters have the largest value, microbiota parameters can also have large values, indicating that both host and microbiota can jointly influence the profiles of these metabolites. Therefore, we decided to investigate metabolites whose profiles are potentially affected by microbiota activity in more detail.

### Linking microbiota-associated metabolites with bacterial activities across conditions

To investigate potential microbiota contributions to the metabolite profiles in the GIT, we inspected microbial metabolism parameters estimated by the intestinal flux model. We defined potential microbial substrates as those for which the normalized microbial metabolism parameter is ≤ −0.5 (a negative value that is at least 50% of the absolute value of the largest parameter for that metabolite), and potential microbial products with microbial metabolism parameters ≥ 0.5. In total, 547 metabolites in the GIT were defined as potential microbial substrates, and 570 as potential microbial products (Figure 5a). To test whether potential substrates and products can be linked by metabolic activity, we searched for enzymatic reactions that connect them in the KEGG database^39^ (Figure 5b). We found 202 potential substrate-product pairs that can be connected by a single enzymatic reaction, and 1,503 pairs and 5,263 pairs that can be connected by two or three subsequent enzymatic reactions, respectively, suggesting that some of the identified substrate-product pairs can reflect microbial metabolic activity in the gut (Supplementary Table 9). Further, the levels of 21 (out of 547) potential substrates and 27 (out of 570) potential products were also significantly different between colonized and germfree mice in serum and/or liver, while in total 20% of systemic metabolites were affected by mouse colonization state (294/1,241 metabolites in serum and 160/1,733 metabolites in liver, FDR ≤ 0.1, Supplementary Table 7).

**Figure 5.**
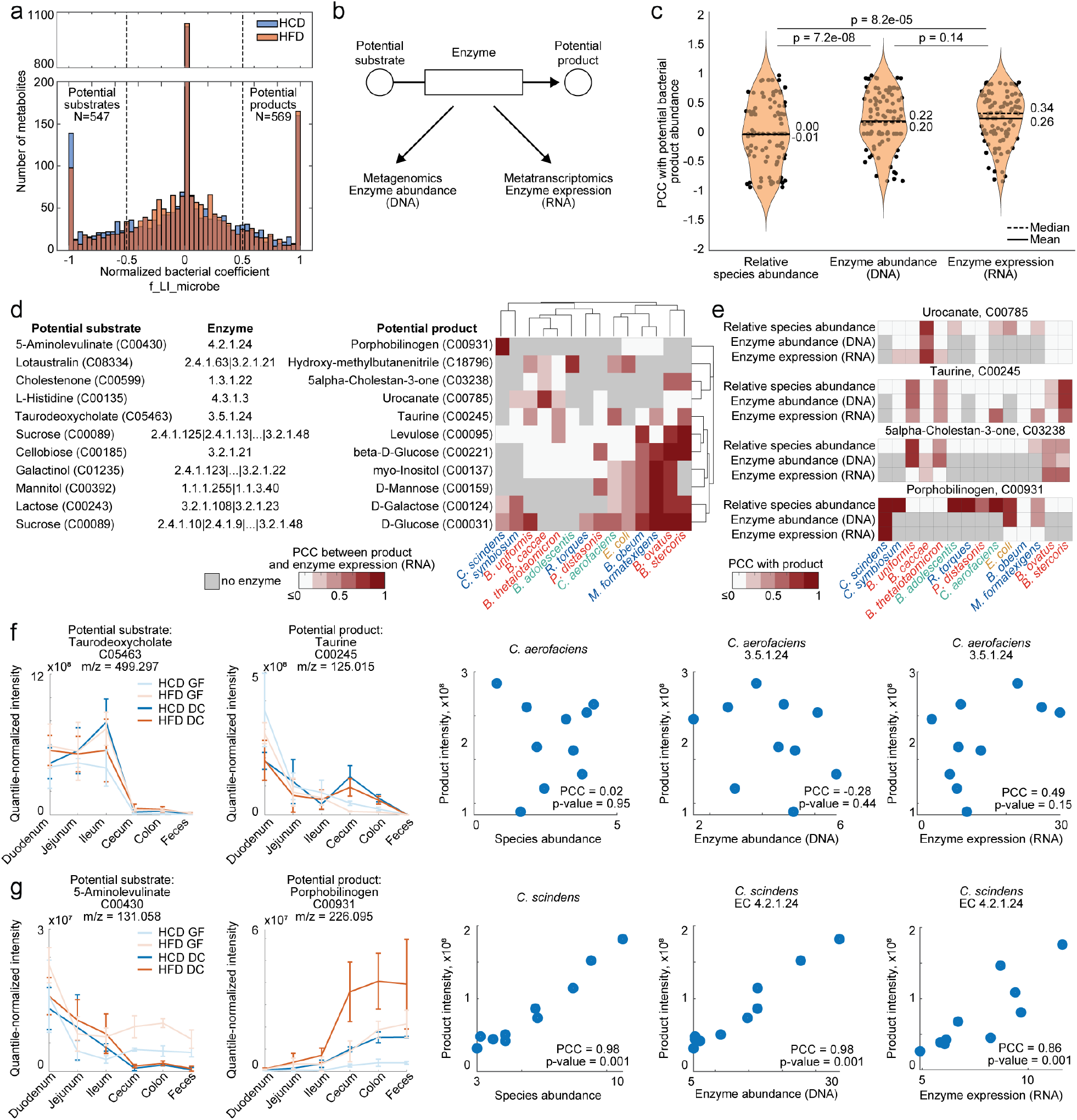
Identifying potential microbial substrates and products and bacterial species and enzymes affecting their levels in the gut. **a.** Distribution of bacterial metabolism coefficients identified by the intestinal metabolic flux model across either of the diet conditions and all metabolites stably detected in the GIT, for which profiles restored in the reverse problem correlated at PCC ≥ 0.7 with the original data. **b.** Schematic representation for potential substrate-product pairs analysis, where enzymatic paths connecting potential substrates to potential products were extracted from the KEGG database, and the corresponding enzyme abundance and expression was calculated from metagenomics and metatranscriptomics data, respectively. **c.** Distributions of PCC between potential bacterial products and either of the three types of microbiota measurements: relative species abundance, gene abundance of the associated KEGG enzyme detected with metagenomics, or expression of the associated KEGG enzyme detected with metatranscriptomics, for all potential products connected by a single enzyme to a potential substrate. P-values correspond to nonparametric two-sided Wilcoxon signed rank test. **d.** Heatmap representing maximum PCC values between product levels and catalysing enzyme expression (RNA) for each species. Only potential products for which either correlation with enzyme abundance (DNA) or enzyme expression (RNA) is significant in at least one species (PCC p-value ≤ 0.1) are depicted. Gray color indicates that there is no catalysing enzyme annotated in the genome, white color indicates PCC ≤ 0.2. **e.** Heatmaps representing PCC values between metabolite levels and relative species abundance, gene abundance (DNA) or enzyme expression (RNA) for each detected species for selected metabolites. **f,g.** Normalized profiles of potential substrates and products connected by a single enzyme, and scatterplots depicting the relationships between potential product levels and either of the three types of microbiota measurements: relative species abundance, gene abundance (DNA), or enzyme expression (RNA) for selected species. P-value indicates significance of PCC (null hypothesis of non-correlation).

To investigate whether the levels of these metabolites can be directly linked to microbial activity, we performed correlation analysis between metabolite levels and relative species abundance, gene abundance detected with metagenomics, and gene expression detected with metatranscriptomics, taking advantage of the natural variation between individual animals. For the potential substrate-product pairs connected by a single enzyme, the correlation between product abundance and enzyme expression was the highest, followed by enzyme gene abundance and relative species abundance (median PCC = 0.34, 0.20 and 0.00, respectively, Figure 5c), suggesting that enzyme expression detected by metatranscriptomics can better explain the observed variation in metabolite levels than metagenomic measurements.

To identify which members of the synthetic community are affecting metabolite levels, we compared the correlations between the identified microbial products and the catalysing enzyme for each species. Among 66 potential products for which a potential substrate and bacterial enzyme is known, enzyme expression of at least one species significantly correlated with the levels of 13% (9/66) products (PCC p-value ≤ 0.1). We observed both general and species-specific correlations between products and catalysing enzymes. For example, enzyme expression of several species across phyla correlate with the abundance of taurine, suggesting that several species express bile salt hydrolases to cleave taurine from bile acids, as previously reported^48^. Similarly, enzyme expression of multiple species correlates with monosaccharide levels, suggesting that many species break down complex sugars to monosaccharides (Figure 5d, e, f). Notably, at the species and gene abundance levels, only representatives of Bacteroidota correlate with these products, indicating that the contribution of other species would be overlooked without metatranscriptomic measurements (Supplementary Figure 6a-c). For certain metabolites, metagenomics and metatranscriptomic analyses agree on the contributing species, such as for urocanate and for 5alpha-Cholestan-3-one (Figure 5e). We also identified opposite cases, in which metatranscriptomic data helps to narrow down the list of potential contributing species. For example, 2-hydroxy-2-methylbutanenitrile, an intermediate of cyanoamino acid metabolism, and porphobilinogen, an intermediate of heme metabolism, are both correlated with the abundance of multiple species and genes encoding the catalysing metabolic enzymes. Metatranscriptomic data, however, highlights that enzyme expression of a single bacterium (*B. adolescentis* and *C. scindens*, respectively) is correlated with the observed metabolite levels (Figure 5e, g, Supplementary Figure 6c,f, Supplementary Table 10). For 2-hydroxy-2-methylbutanenitrile, 5alpha-Cholestan-3-one and porphobilinogen, their levels in serum and/or liver also differed between colonized and germfree mice (Supplementary Figure 6d-h), indicating that microbial activity in the gut can directly affect the circulating host metabolome.

This analysis identifies potential microbial substrates and products in the gut, which constitute 30% of detected metabolites (1,117/3,716), and suggests 202 direct metabolic links between 70 substrates and 66 products. Integrating metagenomics and metatranscriptomics data pinpoints candidate bacterial species and metabolic enzymes whose levels show significant associations with 13% (6/99) of potential microbial products, 50% of which (3/6) are also more abundant in systemic circulation of colonized mice. This approach can be generally applied to highlight metabolites most likely to be impacted by interpersonal differences in microbiota composition both in the GIT and the systems level.

## Discussion

Separating diet, host and microbiome contributions to metabolite levels is a challenging task, especially since host and microbes share many metabolic capacities. Physiology-based pharmacokinetic models provide one strategy to disentangle host and microbiome contributions to metabolism of xenobiotics such as medical drugs^49,50^. However, this approach relies on drug and metabolite profiles collected across mouse tissues over a time-series following drug administration. For metabolites that may originate from diet, host, or microbiome, time-series measurements are not feasible to obtain because host and microbial metabolites are continuously produced. In this study, we found that spatial metabolite distributions across mouse tissues are consistent across different conditions and between individual animals even though the feeding and sample collection times were not synchronized. This suggests that physiological processes in the body are the main determining factors of metabolite distributions in different tissues. This allowed us to assume a pseudo-steady state of metabolite levels in different sections of GIT and develop a coarse-grained intestinal metabolic flux model that describes major factors affecting metabolite abundance. This model estimates the relative contributions of diet, host, and microbiota activity to metabolite profiles by comparing the corresponding parameter values. Although we interpreted metabolites with large negative or positive bacterial metabolism parameters as potential bacterial substrates or products, this approach cannot distinguish microbiome consumption or production from microbiome-dependent changes in the host consumption or production of these metabolites. We demonstrated that combining the modelling approach with other types of data reflecting microbial metabolic activity, such as metagenomics and metatranscriptomics, highlights metabolites that are more likely to be directly metabolized by the microbes.

Metatranscriptomics data has been underutilized in microbiome research due to experimental and computational challenges along with high processing costs^51^. Some studies comparing metagenomics and metatranscriptomics data from the same samples suggest that expression data may not provide much additional value to investigate microbiome function^29,52^. However, while metatranscriptomics and metagenomics estimates of species abundances are generally consistent in our study, we observed large discrepancies between relative RNA and DNA abundances in certain species (Supplementary Figure 2d). While we cannot exclude a technical explanation of this observation such as differences in RNA extraction efficacy or RNA degradation, our data suggests that at least for some species, metatranscriptomic and metagenomic contents of a microbial community can substantially vary. Further, our approach to integrate metatranscriptomics and metabolomics data based on the knowledge of enzymatic reactions connecting potential bacterial substrates and products demonstrated multiple cases in which the species-metabolite correlations produced from metagenomic data (species or gene abundances) were distinct from the correlations produced from metatranscriptomic data (enzyme expression). Notably, correlations of metabolite levels with gene abundances estimated from metagenomic data were distinct from correlations with species abundances, likely reflecting differences in genome replication. These results underline that while levels of some metabolites can be related to microbial species abundance, for others gene abundance is required to explain changes in metabolite levels, while in many cases information about enzyme expression best explains the differences in metabolite levels.

Finally, although our modelling and multi-omics integration framework mainly focussed on metabolites in the GIT, we found that many of these metabolites were detected in systemic circulation and affected by microbiota presence. This modelling approach can be generally applied to disentangle different factors affecting metabolite profiles in the host in other systems with synthetic or native microbial communities and across a range of diets, host genotypes, disease states, and other conditions. Understanding the relative contribution of different factors to metabolite levels will provide crucial information for the design of targeted intervention strategies and combinational therapies to modulate the host metabolic phenotype and improve human health and dietary response.

## Acknowledgements

We thank the members of the Goodman lab and the Bork lab for helpful discussions; C. Kelly for critical reading of the manuscript, and L. Valle, D. Lazo, T. Wu, the Yale West Campus Analytical Core Center and the Yale Center for Genome Analysis for technical assistance.

## Author Contributions

Study and experiment design: MZ-K, MZ, NB-B, ALG. Data analysis: MZ-K. In vitro experiments: MZ-K, MZ. Gnotobiotic animal work: NB-B, MZ-K. Metabolomics measurements: MZ. Metagenomic and metatranscriptomic sequencing: MZ-K. Manuscript writing: MZ-K, ALG. Manuscript editing: MZ-K, ALG, MZ, PB, NB-B. Funding acquisition: MZ-K, PB, ALG. All authors read and approved the manuscript.

## Materials and Methods

### Microbial strains, growth conditions and synthetic community assembly

*Bacteroides thetaiotaomicron* VPI-5482 *tdk*^53^, *Bacteroides caccae* ATCC 43185, *Bacteroides ovatus* ATCC 8483, *Bacteroides uniformis* ATCC 8492, *Phocaeicola vulgatus* ATCC 8482 (formerly *Bacteroides vulgatus), Parabacteroides distasonis* ATCC 8503, *Bacteroides stercoris* ATCC 43183, *Blautia hydrogenotrophica* ATCC BAA-2371, *Blautia obeum* ATCC 29174, *Clostridium scindens* ATCC 35704, *Clostridium symbiosum* ATCC 14940, *Escherichia coli* BW25113, *Eubacterium rectale* ATCC 33656, *Ruminococcus torques* ATCC 27756, *Marvinbryantia formatexigens* DSM14469, *Collinsella aerofaciens* ATCC 25986, *Bifidobacterium adolescentis* ATCC 15703, and *Bifidobacterium longum* CCUG52486 were cultured on brain heart infusion (BHI; Becton Dickinson) agar supplemented with 10% horse blood (Quad Five) anaerobically at 37° C in an anaerobic chamber (Coy Laboratory Products) containing 20% CO_2_, 10% H_2_, and 70% N_2_. Single bacterial colonies were transferred to 10 mL liquid Gut Microbiota Medium (GMM)^54^. As previously suggested^34^, for *M. formatexigens*, liquid GMM was additionally supplemented with formate at 25 mM final concentration. For *B. hydrogenotrophica*, liquid GMM was additionally supplemented with formate at 25 mM final concentration and ammonium sulfate and ammonium chloride ((NH4)SO4 at 0.3 g/L and NH4Cl at 1 g/L). *B. hydrogenotrophica* was grown in liquid culture for 48 h; *R. torques, B. obeum, B. ovatus* were grown in liquid culture for 24 h; *C. aerofaciens, B. stercoris* and *C. scindens* were grown in liquid culture for 14 h, then diluted 1:10 in fresh liquid medium and grown for another 9 h. All other species were grown in liquid culture for 9 h. At the end of incubation, 3 mL of liquid culture of each species was combined, final density of each species was measured with CFU plating, and the synthetic community mixture was stored in 1mL aliquots in 20% glycerol stocks (by mixing 40% glycerol in LB in equal amounts with the culture) in Wheaton vials with anaerobic headspace at −80°C until further use.

### Gnotobiotic Animal Experiments

All experiments using mice were performed using protocols approved by the Yale University Institutional Animal Care and Use Committee in accordance with the highest scientific, humane, and ethical principles and in compliance with federal and state regulations, such as Animal Welfare Act (AWA) and the Public Health Service (PHS).

Germfree (GF) 9-16 week old C57BL/6J mice were maintained in flexible plastic gnotobiotic isolators with a 12-hour light/dark cycle and GF status monitored by PCR and culture-based methods. Before the start of the diet experiment, individually caged animals (n = 5 per group, littermates of mixed sex were randomly assigned to experimental groups) were fed a standard, autoclaved mouse chow (5K67 LabDiet, Purina) *ad libitum*.

### Mouse Colonization Experiments

For the colonization experiment, 1 mL aliquots of synthetic community mixture were thawed on ice, and 200 μL (~10^8^ CFUs) were administered to each animal by oral gavage. On the next day after gavage, standard mouse chow was switched to either irradiated High-fat diet (HFD; TD.06414, Envigo Teklad Diets) or calorie-matched irradiated High-carbohydrate diet (HCD; TD. 140806, Envigo Teklad Diets) *ad libitum*. For the entire duration of the diet experiment, mice were kept in individual cages.

Four weeks after the start of diet administration, mice were euthanized. Liver, blood from the heart, intestinal contents separated by section (duodenum, jejunum, ileum, cecum, colon, feces) were collected. Blood was kept on ice; all other samples were frozen on dry ice immediately. Blood samples were centrifuged for 5 min at 4°C 2500 g, serum was transferred to 1.5 mL Eppendorf tubes, centrifuged again for 5 min, transferred into cryovials and stored at −80°C until further processing.

### Metagenomics and Metatransciptomic Sample Preparation and Sequencing

For DNA and RNA extraction, aliquots of ~100 mg of frozen cecal samples were thawed in 1 mL of RNAProtect, resuspended, incubated at RT for 2 minutes, and pelleted by centrifugation for 1 minute at 15,000 × g at 4°C. Nucleic acids were then purified as described ^34^. Briefly, pellets were resuspended in a solution containing 250 μL of acid-washed glass beads (Sigma-Aldrich), 500 μL of extraction buffer A (200 mM NaCl, 20 mM EDTA), 210 μL of 20% SDS, and 500 μL of phenol:chloroform:isoamyl alcohol (125:24:1, pH 4.5; Ambion), and lysed in a bead beater high for 4 min (2 min / kept on ice 1 min / 2 min) (BioSpec Products). Cellular debris was removed by centrifugation (8,000 × g, 4°C, 5 min). The extraction was repeated, and the nucleic acids were precipitated with isopropanol and sodium acetate (pH 5.5). The samples were placed in a −80°C freezer for 7.5 min, then centrifuged at 12,700 rpm / 4°C / 15 min (~18,200 × g), supernatant was removed, 900 μl cold 100% ethanol was added, the samples were mixed and vortexed briefly, centrifuged at 12,700 rpm / 4°C / 5 min, supernatant was decanted, and the samples were dried in a speed-vac at ambient temperature for 5 min. The pellet was resuspended in 150 uL nuclease-free water, 100 uL of the sample was used for RNA purification, and the remaining 50 uL for DNA purification.

#### RNA purification

The crude extracts were cleaned using the RNeasy Mini kit (Qiagen) with on column DNase I treatment, treated with Baseline-ZERO DNase (Epicentre), and cleaned again by RNeasy Mini. RNA quality was assayed on 2100 Bioanalyzer (Agilent), and only samples with RIN score above 8 were used for metatranscriptomic measurements.

#### DNA purification

50 uL of TE buffer was added to the ~50 uL of the DNA sample in nuclease-free water, followed by adding 0.5 uL RNAse1 and incubation at RT for 2 min. DNA cleanup was performed using a Qiagen PCR purification kit. DNA was eluted in 32 μL EB buffer and quantified on a Nanodrop instrument.

#### Preparation and sequencing of metagenomics and metatranscriptomics libraries

For metatranscriptomics sequencing, rRNA was depleted using the NEBNext rRNA Depletion Kit (NEB). Metagenomics and metatranscriptomics libraries were prepared using Kapa Biosystems Hyper prep kit WGS (Roche) with multiplexing, as recommended by the manufacturer. Metagenomics and metatranscriptomics sequencing wase performed at the Yale Center for Genome Analysis using Illumina NovaSeq6000 sequencer and 2×150 bp sequencing to generate ~10-30 million reads per sample.

### Metabolomics Experiments

#### Extraction of solid tissue samples

200 μL of 0.1 mm zirconia/silica beads (BioSpec Products) and 500 μL of organic solvent (acetonitrile:methanol, 1:1) were added to 50-350 mg of pre-weighed solid tissue material. Material was homogenized by mechanical disruption with a bead beater (BioSpec Products) set for 2 minutes on high setting at room temperature. After incubation for at least 1 h at −20°C, samples were centrifuged (3220 rcf, −9°C) for 15 min. 10 μL of supernatant were diluted with 10 μL H_2_O for analysis by LC-MS.

#### Untargeted metabolomics

Untargeted metabolomics measurements were carried out using an Agilent 1290 UHPLC system coupled to an Agilent 6550 qTOF mass spectrometer. Chromatographic separation was performed on either of the three columns: a ZIC-pHILIC column (Merck, 150 mm × 2.1 mm, 5 μm particle size), a reverse-phase C18 Kinetex Evo column (Phenomenex, 100 mm x 2.1 mm, 1.7 μm particle size), or a reverse-phase C18 Kinetex Evo column (Phenomenex, 100 mm x 2.1 mm, 1.7 μm particle size). Chromatographic separation with ZIC-pHILIC column was performed using mobile phase A: 10 mM ammonium carbonate buffer (pH 9.3) and B: acetonitrile;. 3 μL of sample were injected at 80% B and 0.1 mL min^-1^ flow followed by a linear gradient to 50% B over 20 min and 0.1 mL min^-1^ flow ^55^. The column was re-equilibrated at starting conditions for 8 min. Chromatographic separation with C8 and C18 columns was performed using mobile phase A: H2O, 0.1% formic acid and B: methanol, 0.1% formic acid at 45°C. 5 μL of sample were injected at 100% A and 0.4 mL/min flow followed by a linear gradient to 95% B over 5.5 min and 0.4 mL/min. The qTOF was operated in positive (50 - 1000 m/z) and negative (50 - 1050 m/z) scanning mode and source parameters optimized following the manufacturer’s recommendations: VCap: 3500 V, nozzle voltage: 2000 V, gas temp: 225°C; drying gas 13 L min^-1^; nebulizer: 20 psg; sheath gas temp 225°C; sheath gas flow 12 L min^-1^. Online mass calibration was performed using a second ionization source and a constant flow (5 μL min^-1^) of reference solution (positive mode: 121.0509 and 922.0098 m/z and negative mode: 112.9856 and 1033.9881 m/z). The MassHunter Quantitative Analysis Software (Agilent, version 6.0) was used for peak integration.

### Metagenomics and metatranscriptomics sequencing analysis

#### Bacterial abundance analysis

Species abundance analysis from metagenomics and metatranscriptomics samples was performed with MetaPhlAn 2^56^. For further analysis, bacterial species (filtered by taxonomic level “|s”, “unclassified” species were filtered out) which were detected in at least 5 samples with abundance above 0.01% were selected. For abundance clustergrams, relative abundance of each species was normalized across all samples with zscore function in MatLab 2019a, and clustergrams were built with clustergram function in Matlab 2019a. Species abundances between mouse groups were compared with nonparametric two-sided Wilcoxon-Mann-Whitney test (ranksum function in Matlab 2019a).

#### Gene and transcript abundance analysis

Reference genomes and annotations for single species were downloaded from NCBI (Supplementary Table 1) in fasta and gff format and additionally annotated using EggNOG-Mapper^57^. Metagenomics and metatranscriptomics raw reads were preprocessed with KneadData^58^ to trim sequencing adapters with Trimmomatic^59^ v0.38 (--trimmomatic-options =”ILLUMINACLIP:TruSeq3-PE.fa:2:30:10” --trimmomatic-options=”LEADING:3” --trimmomatic-options=”TRAILING:3” --trimmomatic-options=”SLIDINGWINDOW:4:20” --trimmomatic-options=”MINLEN:36”) and remove mouse reads with built-in database mouse_C57BL (-db mouse_C57BL). Read quality was assessed with FastQC v0.11.8. Reference genomes were prepared for HISAT2^60^ analysis by building HISAT2 index from downloaded gff files with hisat2_build function. Preprocessed reads sequences were aligned to the reference genomes with HISAT2, sam files were converted to bam files with samtools^61^, transcript abundances were estimated with StringTie v2.0^62,63^ and prepared for further analysis with prepDE script provided together with StringTie. Differential expression analysis between the mouse groups was performed with EDGE-R^64^ v3.28.1, and reads were normalized using GetMM^65^ in RStudio (R v3.6.1).

#### Genome coverage analysis

Geneome coverage by metagenomics and metatranscriptomics was defined as the number of genes/transcripts that had a count higher than 0, divided by the total number of predicted genes per species in annotations downloaded from NCBI (Supplementary Table 1). For comparison of MetaPhlAn estimates species abundances with species abundance estimated from metagenomics and metatranscriptomics, abundance estimates were produced by counting the number of raw reads mapping to each genome included in the study and dividing by the total number of reads per sample.

#### Estimation of gene abundance and enzyme expression

Enzyme annotation was obtained from the EggNOG annotation tables. For enzymes of interest, their gene abundance and enzyme expression per species was calculated as summed GetMM-normalized gene (metagenomics) or enzyme (metatranscriptomics) counts for all genes annotated as the specified enzyme. Pearson’s correlation coefficients were calculated between summed gene abundances and enzyme expression values and quantile-normalized metabolite abundances summed for cecum, colon and feces. For each potential microbial product for which several catalysing enzymes exist, the enzyme with the largest positive correlation was kept for plotting.

### Metabolomic data filtering and annotation

Intensities of ions detected by each method in each tissue were first normalized by tissue weight, and ions that were detected in <5 samples per tissue were filtered out. Per method, ions detected in different tissues were combined in one table based on ion mass over charge ratio (m/z), retention time (RT), and detected spectrum. Ion detected across methods and tissues were combined in one table for further analysis. Ions were annotated to metabolites based on exact mass considering [M-H^+^] and [M+H^+^] ions using the metabolite reference list compiled from the Kyoto Encyclopedia of Genes and Genomes (KEGG) metabolite repository^39^ and Human Metabolome Database (HMDB)^40^. Ions were assigned to metabolites allowing a mass tolerance of 1 mDa and an intensity cut-off of 5,000 counts. Tissue weight-normalized intensity values were further quantile-normalized with quantnorm function in MatLab 2019a (MathWorks). Annotated metabolites were scored based on RT values, intensity correlation across methods, intensity value across samples, number of samples where metabolite was detected, and number of carbon isotopes detected in the ion spectrum. For each metabolite, only the annotation with the top score was retained.

### Statistical data analysis and spatial clustering

#### Principal component analysis

Metabolomics, metagenomics and metatranscriptomics data were imported into MatLab 2019a (MathWorks) for quantitative and statistical analysis. Principal component analysis was performed on log_10_-transformed quantile-normalized metabolomics data, or log_10_-transformed GetMM-normalized metagenomics and metatranscriptomics data with pca function in MatLab.

#### Differential metabolite abundance analysis

For differential analysis of metabolomics data, quantile normalized metabolite intensities were compared between mouse groups in each diet condition, and between diet conditions within each mouse group with two-sided t-test with equal variances (ttest2 function in MatLab). P-values were corrected for multiple hypotheses testing by calculating the false-discovery rate (FDR) with the Benjamini-Hochberg procedure (mafdr function in MatLab).

#### Spatial metabolite profile clustering

For spatial metabolite profile analysis, per each mouse group and dietary condition, mean metabolite intensity per GIT tissue was calculated, and mean intensities across the six GIT sections were normalized with zscore function in MatLab. To select the optimal k for k-means cluster analysis, silhouette criterion was used in evalclusters function in MatLab (with parameters (‘kmeans’,‘silhouette’, ‘KList’, 1:30) to test k values between 1 and 30) for each of the four conditions separately. To test robustness of cluster assignment, k-means clustering procedure for the optimal k (k=6 for each group) was performed 100 times, and final cluster assignment for each metabolite was chosen based on the most frequent cluster assignment. For serum and liver, three clusters were defined based on metabolite detection across tissues: Cluster 1 detected only in serum (or only in liver), Cluster 2 detected in serum and liver (or liver and serum), and Cluster 3 detected in serum and GIT (or liver and GIT).

### Pathway and chemical group enrichment analysis

For pathway enrichment analysis, KEGG pathway-gene assignments were extracted from the EggNOG annotation tables, and only pathways annotated in at least one species were analysed. For chemical group enrichment analysis for metabolites, chemical group information was downloaded from HMDB database^40^. Enrichment procedure based on the gene set enrichment analysis (GSEA) method^66^ was applied as described^67^. Significantly changing transcripts (|log_2_(fold change)| ≥ 1, FDR ≤ 0.05) were ranked either by fold change or by FDR, and enrichment p-values were calculated with the Fisher exact test for each subset of size varying from 1 to the total changing set size. For each pathway, the smallest p-value of all the subsets was retained, and p-values were adjusted for multiple hypotheses testing by calculating FDR with the Benjamini-Hochberg procedure. Chemical group enrichment analysis of metabolites belonging to different k-means clusters or different groups defined from hierarchical clustering of the model coefficients was performed in the same way without varying the set size.

### Substrate-product enzyme path analysis

For KEGG substrate-product-enzyme network analysis, the KEGG reaction pair list, KEGG reaction pair, and KEGG compound list were downloaded from KEGG API (application programming interface) (http://rest.kegg.jp/, accessed July 2015), and only the main (substrate-product) reaction pairs were retained. These datasets were used to define the metabolite adjacency matrix, where each metabolite is connected to another one if they are substrate and product in a reaction. K-shortest path script based on Yen’s algorithm^68^ (implementation by Meral Shirazipour (2011) K-Shortest Path - Yen’s algorithm http://www.mathworks.com/matlabcentral/fileexchange/32513-k-shortest-path-yen-s-algorithm, accessed July 2015) was used to calculate the shortest paths between each two metabolites based on the connectivity matrix. Information on enzymes associated with each reaction was downloaded from KEGG API (https://rest.kegg.jp/link/reaction/enzyme).

### Intestinal flux model

#### Model overview

The multi-compartment intestinal flux model of metabolite propagation along the GIT contained 5 main compartments (jejunum, ileum, cecum, colon, feces). Metabolite flow from duodenum was included in the model, but duodenum was not explicitly modelled due to the lacking information on the input flux. Metabolite propagation through the GIT was driven by the flow of GI material in different GIT sections following mass action kinetics with condition-independent parameter f. Metabolite levels in jejunum and ileum were additionally affected by the host metabolism flux f_SI_host, which was diet dependent. Metabolite levels in cecum and colon were additionally affected by the host metabolism flux f_Cecum_host and f_Colon_host, which were diet dependent. In colonized mice, metabolite levels in cecum were additionally affected by microbial flux parameter f_LI_microbe, also diet-dependent. Model parameters and equations are provided in Supplementary Table 8.

#### Forward problem

With the pseudo-steady state assumption, 20 model equations (for each of the 5 compartments for each of the 4 mouse groups) represented a system of linear equations in respect to 9 parameters (f, f_SI_host_HCD, f_Cecum_host_HCD, f_Colon_host_HCD, f_LI_microbe_HCD, f_SI_host_HFD, f_Cecum_host_HFD, f_Colon_ost_HFD, f_LI_microbe_HFD). For the parameter fitting, metabolite intensities (that were coefficients of the intestinal flux f) were normalized to the maximum value across the 5 tissues in all 4 mouse groups. The model parameters were identified by solving a linear optimization problem with lsqlin function in MatLab 2019b (MathWorks) with 0 as a lower limit for intestinal flux parameter f with no upper limits, and no lower or upper limits for the other parameters.

#### Reverse problem

To solve the reverse problem, the same system of linear equations was rewritten to represent flux values identified in the forward problem as coefficients, and metabolite intensities as parameters that need to be estimated. The model parameters were identified by solving a linear optimization problem with lsqlin function in MatLab 2019b (MathWorks) with 0 as a lower limit for all parameters representing metabolite levels and no upper limits.

## Data and code availability

The raw metagenomics and metatranscriptomics data have been deposited in the EMBL-EBI European Nucleotide Archive under accession no. PRJEB33195. Processed data are provided in Supplementary Table 4. The metabolomics data are provided in Supplementary Tables 6-7. Supplementary Tables are available on Zenodo (https://doi.org/10.5281/zenodo.6991165). Analysis pipelines and the modelling framework are available at https://github.com/mszimmermann/diet_microbiota_host_git_model.

## Supplementary Materials

### List of Supplementary Tables

**Supplementary Table 1.** List of 18 genome-sequenced human gut bacteria with metabolic characteristics that were used for community assembly.

**Supplementary Table 2.** Diet composition

**Supplementary Table 3.** Species relative abundance

**Supplementary Table 4.** Gene abundance and expression changes

**Supplementary Table 5.** Gene pathway enrichment results

**Supplementary Table 6.** Metabolomics data

**Supplementary Table 7.** Metabolite fold changes, clustering and model parameters

**Supplementary Table 8.** Description of the intestinal flux model

**Supplementary Table 9.** Enzymatic paths between substrates and products

**Supplementary Table 10.** Pearson’s correlation coefficients between potential substrates and products, and metagenomics and metatranscriptomic measurements

### List of Supplementary Figures

**Supplementary Figure 1.**
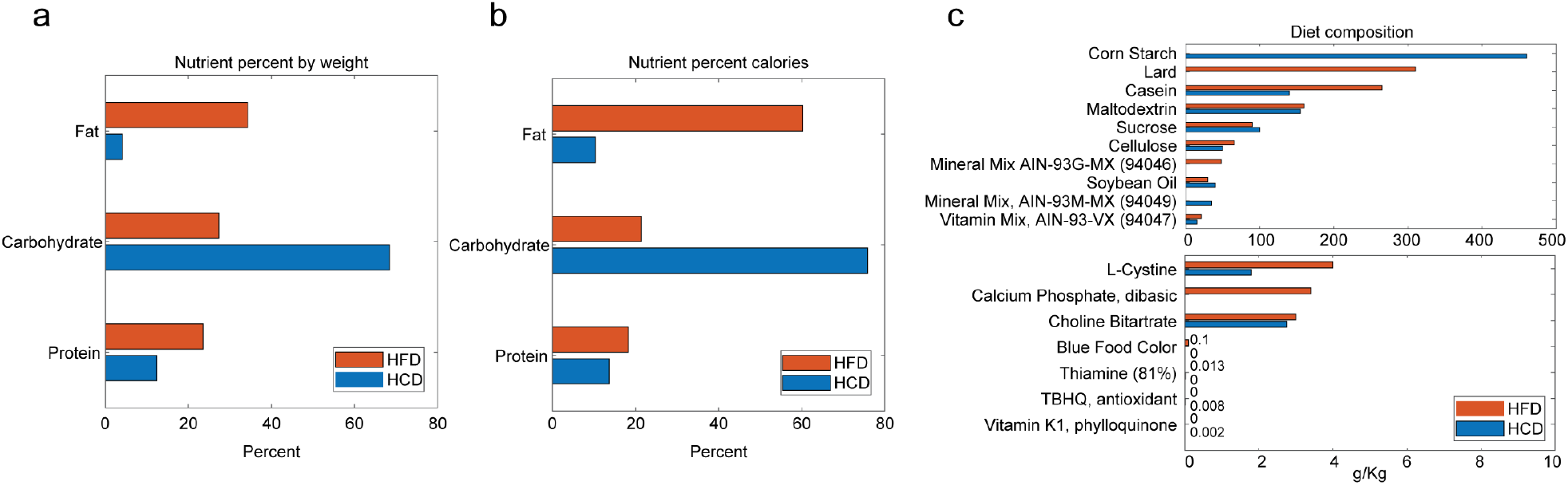
Composition of HCD and HFD diets. **a.** Nutrient percent by weight in each of the two diets. **b.** Nutrient percent by calories in each of the two diets. **c.** Detailed nutrient composition in each of the two diets. HFD – high-fat diet, HCD – high-carbohydrate diet.

**Supplementary Figure 2.**
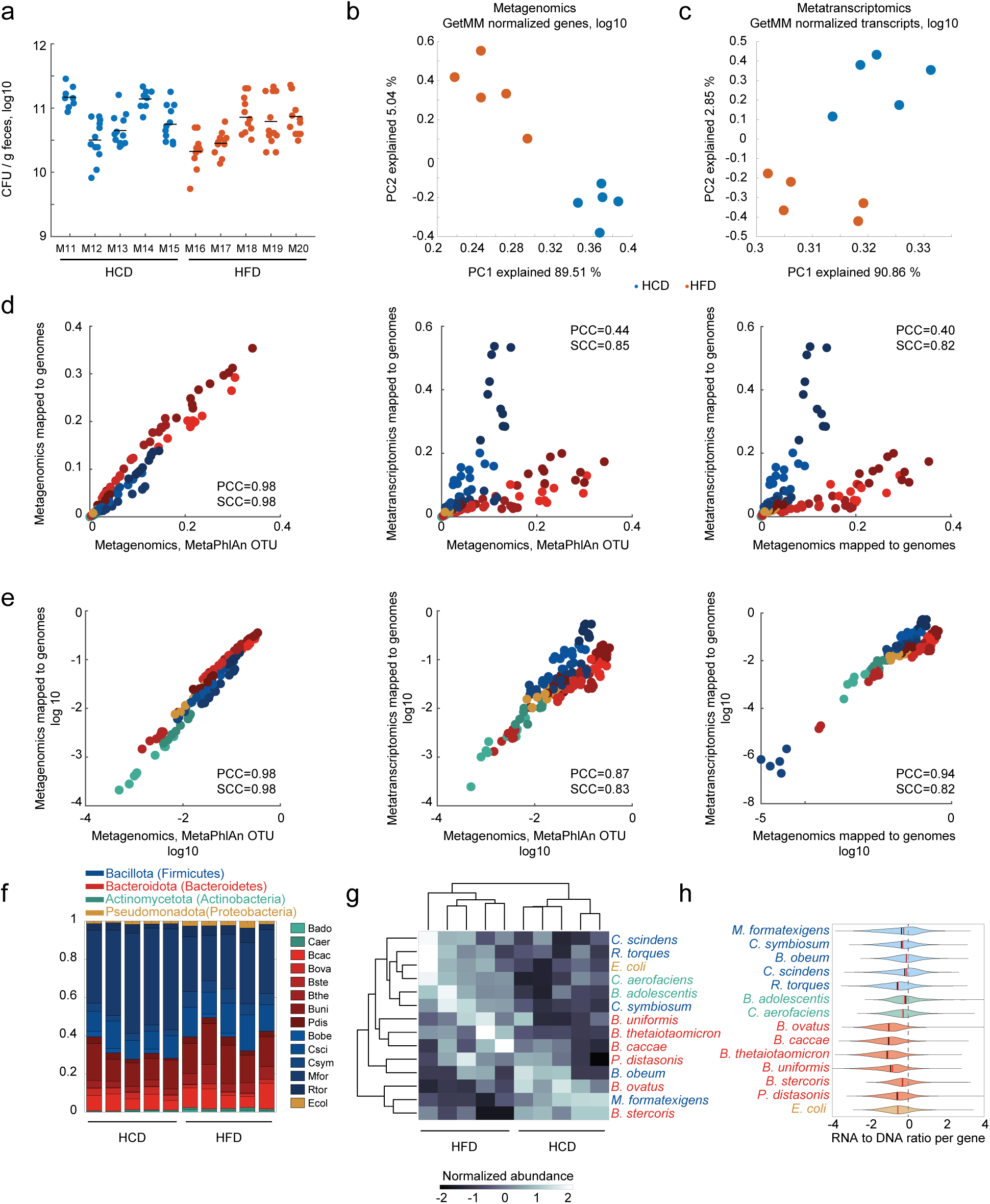
Concordance and differences between metagenomics and metatranscriptomics measurements of the mouse microbiota. **a.** Colony forming units (CFU) counts of bacteria per gram of mouse feces at the end of experiment. Individual animals are depicted m11-m20, each dot within each animal represents different dilutions of technical replicates for the fecal cultures. **b.** PCA of log10-transformed metagenomics measurements normalized in the GetMM pipeline. **c.** PCA of log10-transformed metatranscriptomics measurements normalized in the GetMM pipeline. **d.** Comparison of species abundance estimate between MetaPhlAn pipeline (metagenomics) and species estimate from mapping metagenomic data to the genomes of community members (left); MetaPhlAn and metatranscriptomics mapped to individual genomes (center), metagenomics and metatranscriptomics (right). Each point represents the relative abundance of a species in an individual animal. **e.** The same as (d) where species abundances are represented in log10 scale. **f.** Stacked barplot representing relative species abundance estimated from metatranscriptomics data across individual animals. **g.** Hierarchical clustering of normalized species abundances in (f). **h.** Distributions of RNA to DNA ratios calculated from metatranscriptomics and metagenomics measurements per gene for each species across all samples. PCC – Pearson’s correlation coefficient, SCC – Spearman correlation coefficient.

**Supplementary Figure 3.**
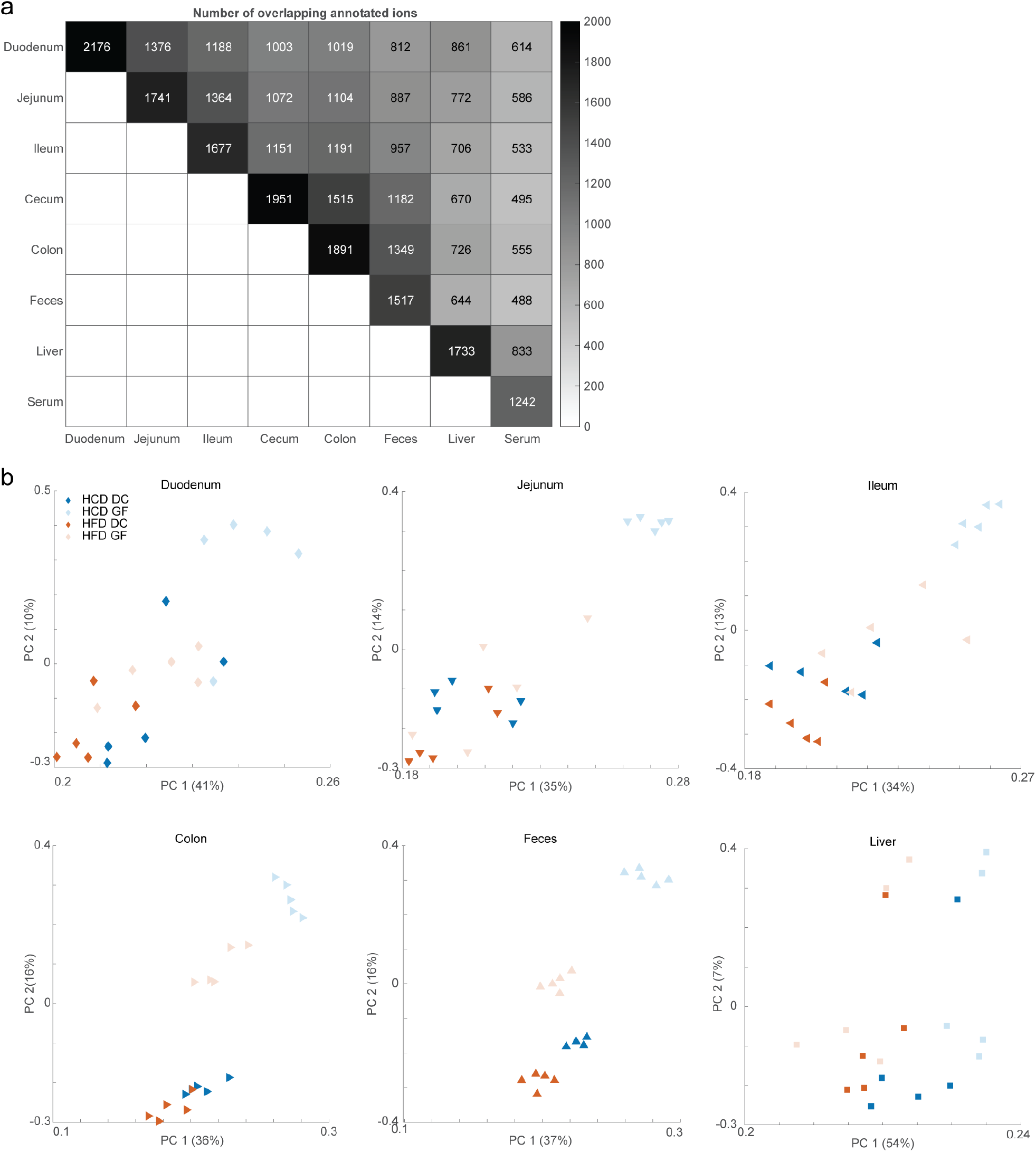
Comparison of metabolomic measurements across tissues. **a.** Number of annotated compounds measured in each tissue (diagonal) and overlapping between each pair of tissues (in row and column). **b.** PCA of annotated metabolites in individual tissues.

**Supplementary Figure 4.**
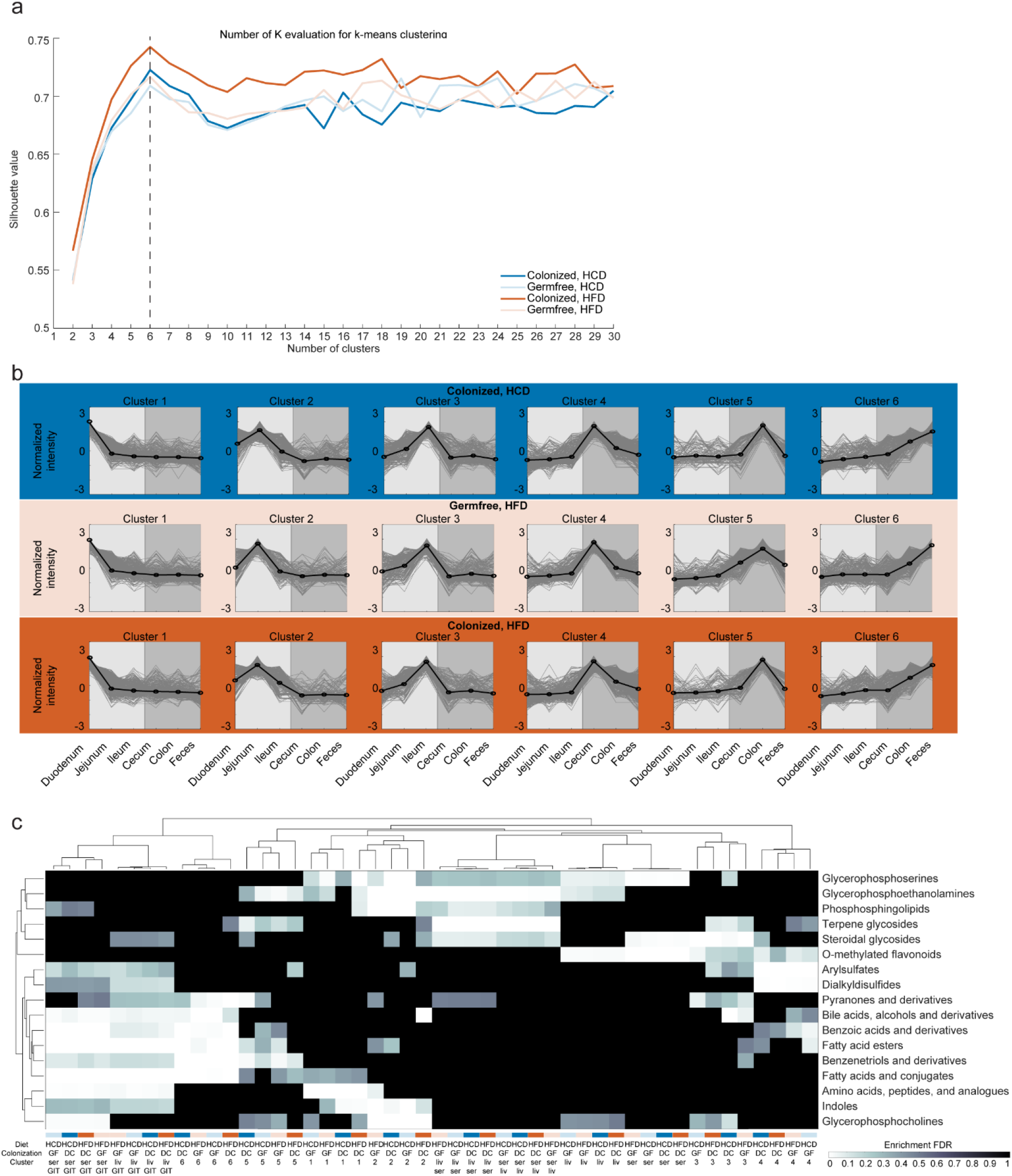
Clustering annotated metabolites according to spatial distribution across tissues. **a.** Silhouette criterion values evaluate clustering of GIT metabolite profiles for different number of k-means clusters (between 2 and 30) for each of the four mouse groups separately. The cluster number resulting in the optimal silhouette criterion value (the highest) is marked with a dashed line. **b.** K-means clusters of normalized metabolite profiles in the GIT of germfree mice consuming HFD and colonized mice consuming either HCD or HFD. **c.** Hierarchical clustering of the results of chemical class enrichment analysis of the metabolites belonging to each of the k-means clusters combined with clusters of metabolites detected in serum or liver only, overlapping between serum and liver, or present in serum and GIT or liver and GIT. Analysis was performed separately for each cluster in each of the four mouse groups. Only significantly enriched groups (FDR-adjusted p-value ≤ 0.1) are depicted. HFD – high-fat diet, HCD – high-carbohydrate diet, GF – germ-free mice, DC – mice colonized with the defined community.

**Supplementary Figure 5.**
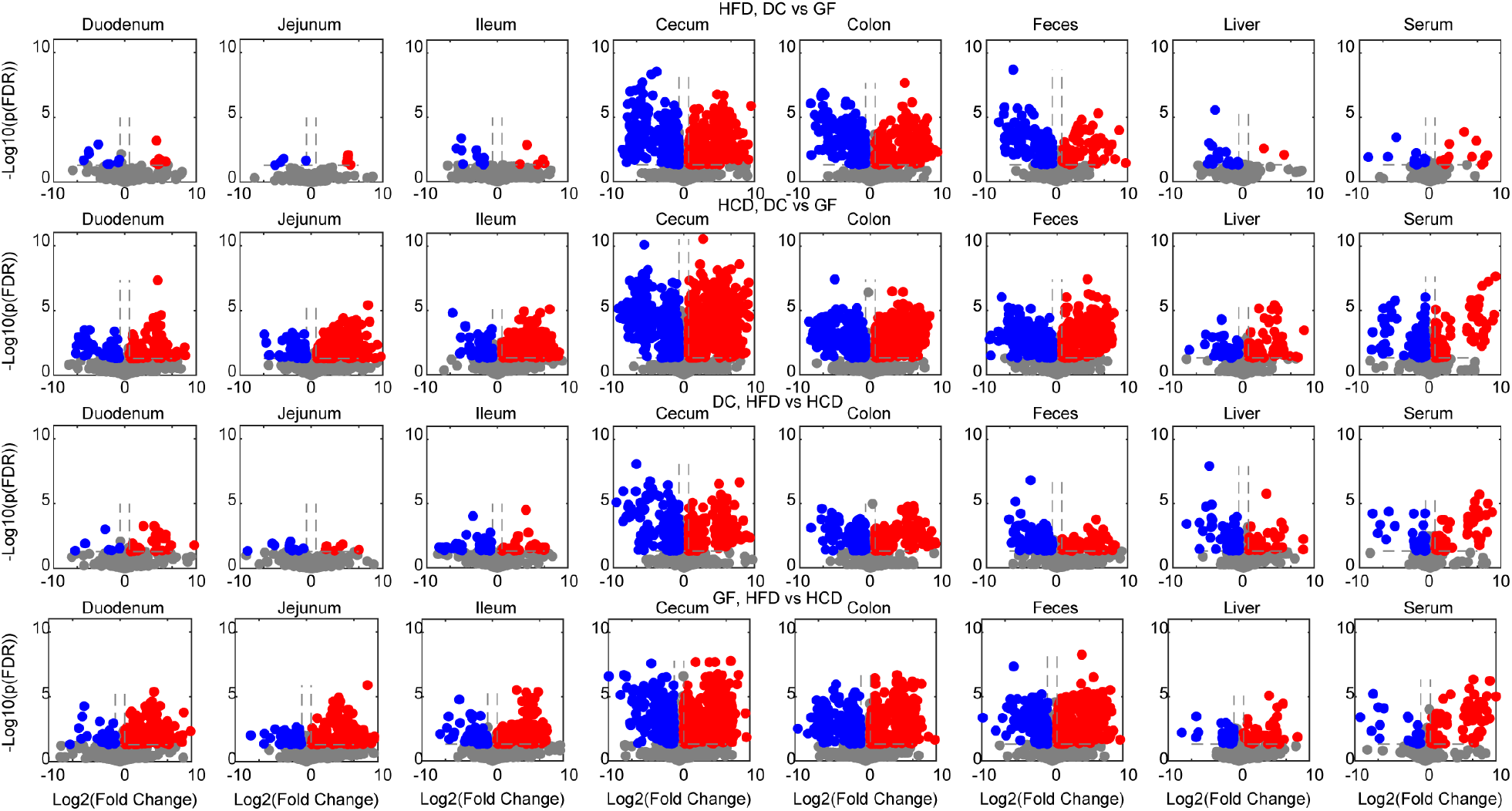
Differential metabolite abundance across tissues, diets and mouse groups. Volcano plots depicting significantly differentially abundant metabolites ((abs(log2(fold change)) ≥ 1, FDR-adjusted p-value ≤ 0.05)), pairwise comparisons were performed between mouse groups within each diet, or between diets within each mouse group. HFD – high-fat diet, HCD – high-carbohydrate diet, GF – germ-free mice, DC – mice colonized with the defined community.

**Supplementary Figure 6.**
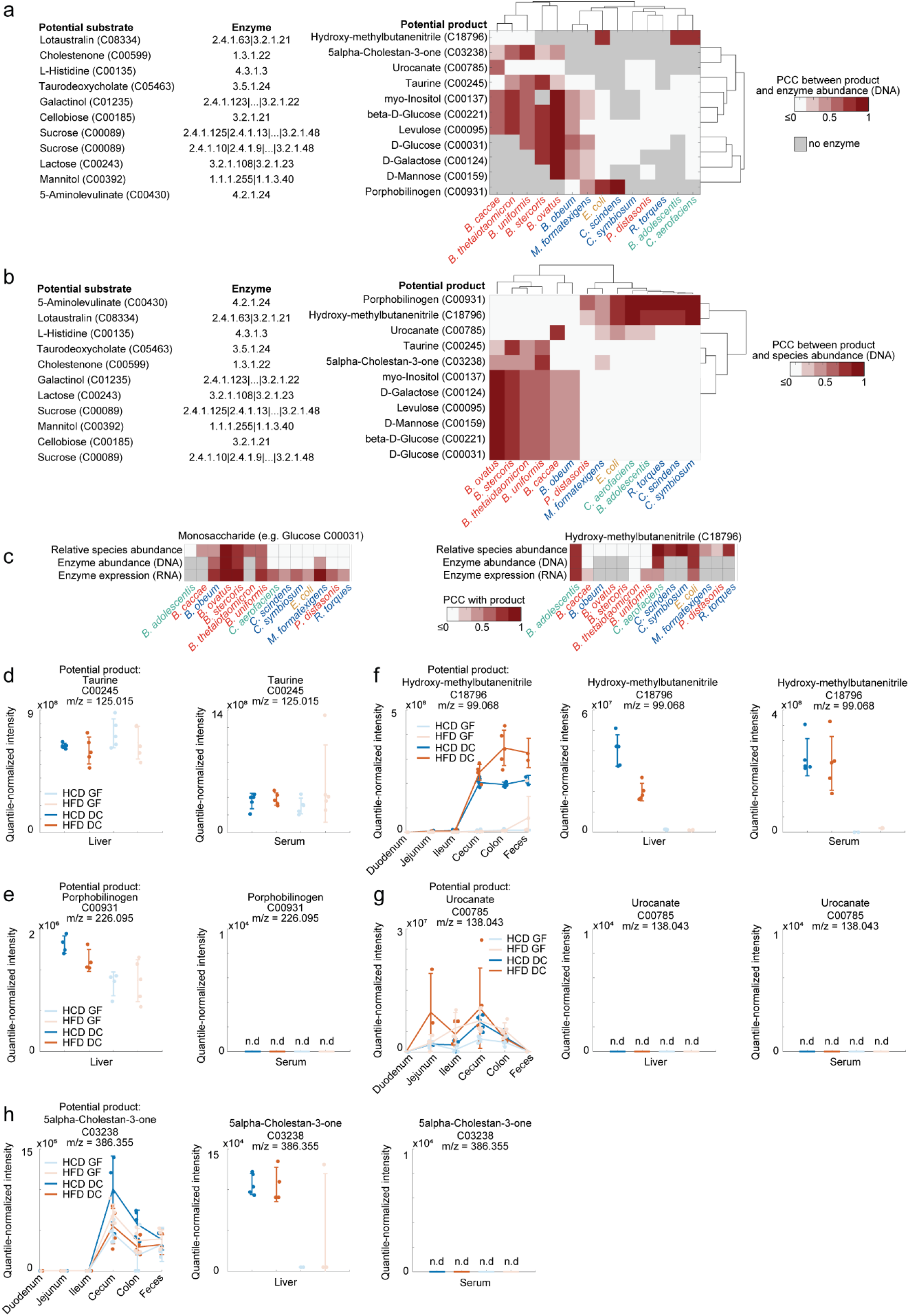
Identifying bacterial species and enzymes affecting potential microbial products in the gut with different types of microbiome data. **a.** Heatmap representing maximum PCC values between product levels and catalysing enzyme abundance (DNA) for each species. Only potential products for which either correlation with enzyme abundance (DNA) or enzyme expression (RNA) is significant in at least one species (PCC p-value ≤ 0.1) are depicted. Gray color indicates that there is no catalysing enzyme annotated in the genome, white color indicates PCC ≤ 0.2. **b.** Heatmap representing maximum PCC values between product levels and species abundance (metaphlan, DNA) for each species. Only potential products for which either correlation with enzyme abundance (DNA) or enzyme expression (RNA) is significant in at least one species (PCC p-value ≤ 0.1) are depicted. White color indicates PCC ≤ 0.2. **c.** Heatmaps representing PCC values between metabolite levels and relative species abundance, gene abundance (DNA) or enzyme expression (RNA) for each detected species for selected metabolites. **d-e.** Quantile-normalized metabolite abundance in liver and serum for potential microbial products depicted in Figure 5. **f-g.** Quantile-normalized profiles of potential products connected by a single enzyme to the corresponding potential substrate, and their levels in serum and liver.

